# Identification and evaluation of besifloxacin as repurposed antifungal drug in combination with fluconazole against *Candida albicans*

**DOI:** 10.1101/2025.03.12.642842

**Authors:** Shuvechha Chakraborty, Ameya Pawaskar, Siddhanath Metkari, Taruna Madan, Susan Idicula-Thomas

## Abstract

Emergence of life-threatening fungal infections like systemic candidiasis concurrently with bacterial infections and limitations of current antifungal therapies warrants the discovery of novel inhibitors. We identified besifloxacin (BS), an FDA-approved antibacterial, as a potent antifungal inhibitor. A combination of besifloxacin with fluconazole showed a positive synergy (δ = 29.58) resulting in 80% inhibition of microbial growth. BS was able to reduce the MIC of FLC from 2mg/L to 0.5 mg/L when used in combination. Additionally in murine systemic *Candida* infection, BS reduced fungal load by 83% in mice kidneys at a dose of 100 mg/kg/day. The findings demonstrated the antifungal potential of BS, proposing its use in combination therapy with fluconazole to combat resistance through alternative mechanisms.

**GRAPHICAL ABSTRACT:** 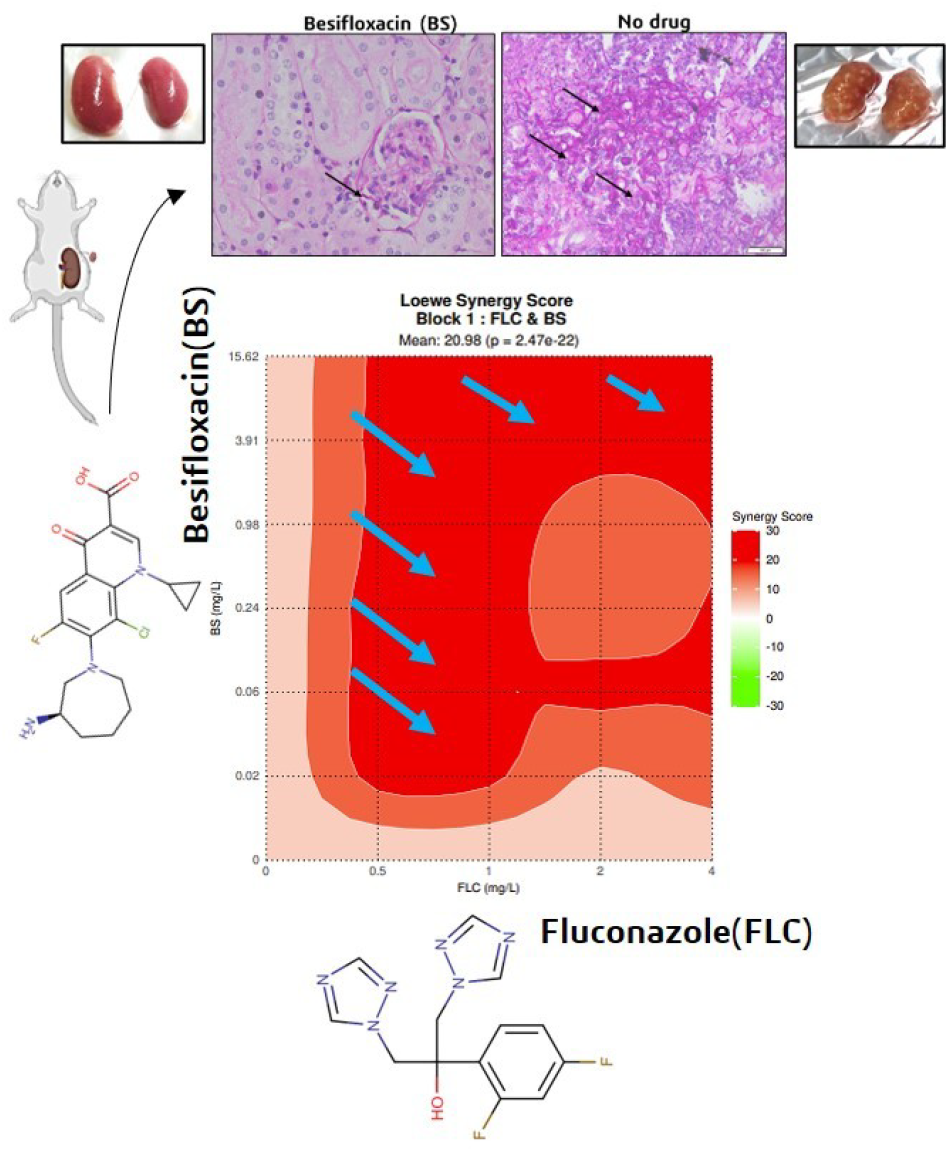

## INTRODUCTION

Invasive fungal infections are typically severe and are associated with individuals who experience mild to severe immunocompromised states due to concomitant health issues. Immunocompromised cohorts are primarily vulnerable to opportunistic pathogens such as *Candida*, which can affect both immunocompetent and immunocompromised hosts, with the latter facing higher vulnerability due to weakened immunity resulting in more severe manifestations of candidiasis (Rayens, Norris, and Cordero 2022).

Most mucosal and systemic infections are caused by *C. albicans*, although other species such as *C. glabrata, C. tropicalis, C. parapsilosis, C. krusei*, and *C. auris* may also be implicated in both forms of candidiasis. While *Candida* spp. is the leading cause of opportunistic mycoses occupying critical to medium priority status in the WHO fungal priority pathogen list 2022 (World Health Organization (WHO) n.d.), the therapeutic options are limited to few antifungal classes. These include azoles, echinocandins, polyenes, and nucleoside analogs. Azoles such as fluconazole, voriconazole, itraconazole are the most frequently used antimycotics for mucosal infections that target fungal lanosterol demethylase enzyme in ergosterol biosynthesis. For severe forms of systemic candidiasis, intravenous antimycotics like polyenes (amphotericin B, nystatin) and echinocandins (caspofungin, micafungin) are the preferred choice; these are often associated with organ toxicity and other undesired side effects (Robbins, Wright, and Cowen 2016). Nucleoside analog fluoropyrimidine that inhibits DNA synthesis is preferred mostly as an adjunct therapy to amphotericin B or azoles due to its fungistatic nature and development of resistance.

The research and development of new antifungal drugs has predominantly focused on the azole and echinocandin classes. This has resulted in a prioritization of reformulations to improve efficacy and safety, while some vaccine candidates (PEV7 and NDV-3) are progressing towards clinical trials (Kaur et al. 2023). The need for new and effective antifungal treatments is of paramount importance, given the increasing prevalence of fungal infections and the emergence of drug-resistant strains. Drug combination therapy is often required alongside azole antifungals in the treatment of candidiasis due to several limitations of azole monotherapy, including resistance development, biofilm formation, and insufficient activity against certain *Candida* spp (Kane and Carter 2022; Lu et al. 2018).

Drug repositioning represents a promising approach that can be employed to enhance drug efficacy, mitigate toxicity, and tackle the growing challenge of drug resistance of currently used antifungals. Notably, drug repurposing can help identify new uses for drugs that have already been approved for clinical use, thereby avoiding the need for extensive preclinical and clinical testing for developing new antifungal treatments.

Multiple studies have indicated that certain antibacterial drugs, including tobramycin, gentamicin, azithromycin, levofloxacin, and tetracycline, exhibit broad-spectrum antifungal properties when evaluated *in vitro* and in animal models. Polymyxin B has been found to impair the permeability of the fungal cell membrane in *Candida albicans* when administered in conjunction with antifungal agents such as ketoconazole, micafungin, and amphotericin B. It binds to anionic lipids present on the fungal membrane with a MIC range of 8 - 256 mg/L, and in combination with fluconazole it compromises the membrane integrity of fungi (Venturini et al. 2016; Zhai et al. 2010). Animal models have also been used to validate the antifungal efficacy of antibacterial drugs. For instance, in the murine candidiasis model, the combination of cefoperazone-sulbactam, colistin, or meropenem with caspofungin resulted in a lower fungal load in the kidneys than caspofungin monotherapy (Keçeli et al. 2014; Keceli Ozcan et al. 2006; Zeidler et al. 2013). Histopathological observations have also indicated that the level of inflammation was lower in the caspofungin and meropenem group than in those treated with caspofungin alone (Keceli Ozcan et al. 2006). Quinolones, when combined with fluconazole and amphotericin B have demonstrated a synergistic effect against invasive candidiasis in mice models (Sasaki et al. 2000; Sugar, Liu, and Chen 1997).

These findings are significant, as clinical studies indicate that around 20-40% of patients with candidemia also experience concurrent bacterial infections(Allison et al. 2016; Chen, Xu, and Wu 2020). In these instances, a broad-spectrum drug with combined antibacterial and antifungal properties in combination with fluconazole could significantly enhance treatment outcomes and lower the risk of azole resistance. To accomplish this goal, we implemented a drug repurposing pipeline to identify broad-spectrum drugs effective against *Candida albicans* (CAL). The selected drug was validated through *in vitro* assays and *in vivo* testing with murine models of systemic candidiasis.

## METHODOLOGY

### Screening for homologs in *Candida* proteome

The reference proteome for CAL strain SC5314 was retrieved from UniProtKB proteomes database. The CD-HIT tool (Huang et al. 2010) was used to obtain a non-redundant proteome of 6022 protein sequences by excluding paralogous sequences. Drug targets from the *Candida* proteome homologous to 70 shortlisted targets(Chakraborty et al. 2023a) were identified by local BLASTp algorithm with e-value cut-off <0.0001. Those targets that exhibited significant alignment with *CAL* proteome were queried against Conserved Domain Database (CDD) using DELTA-BLASTp (Boratyn et al. 2012) to identify homologs with conserved domains in *CAL*. A criteria of query coverage > 80% and e-value cut-off <0.005 was applied to identify sequences with conserved domains. Drugs targeting these conserved domains were used for further validation *in vitro*.

### *In vitro* evaluation of besifloxacin

Stock solutions of the drug besifloxacin cat. no SML1608 and gemifloxacin cat. no SML1625 (Sigma-Aldrich, USA), was prepared in DMSO (dimethyl sulfoxide, cell-culture grade) (Sigma-Aldrich, USA). The stock solution was further diluted in RPMI-1640 media (HiMedia) media supplemented with MOPS (3-(N-morpholino) propanesulfonic acid) and 2% glucose, for assay. For *in vitro* use, the final drug concentration was 1000 mg/L with DMSO not surpassing 0.5% (v/v). Serial dilutions of the drug solution were made in 2X RPMI-1640 in a 96-well plate till the lowest concentration of 0.03 mg/L was achieved. Five colonies of *C. albicans* ATCC MYA-2876 were inoculated in YPD (Yeast Peptone Dextrose) broth overnight at 37 °C in a shaker incubator. Post overnight incubation, the cultures were centrifuged at 4000 rpm for 10 mins, washed, diluted in sterile PBS (phosphate-buffered saline), and adjusted to a cell density of 1 × 10^5^ cfu/mL. An inoculum of 50 μL of the diluted yeast culture was added to each well to evaluate antimicrobial activity. After 24 h incubation at 37 °C, cell viability was checked using XTT ((2,3-bis (2-methoxy-4-nitro-5-sulfophenyl)-5-[(phenylamino)carbonyl]-2Htetrazolium hydroxide) assay kit (Roche) following manufacturer’s protocol.

To gain valuable insights into the potential synergy between besifloxacin and fluconazole in combating *Candida* infections, the anti-*Candida* activity of besifloxacin was evaluated in the presence of fluconazole. Fluconazole concentrations ranging from 4 to 0.5 mg/L were used in combination with besifloxacin at concentrations ranging from 15 mg/L to 0.02 mg/L.

Anti-biofilm assay [Wong et al., 2014] was performed to assess the effect of the drug besifloxacin on the *CAL* biofilms. Briefly, 100 μL of 10^6^ cells /mL in YNB media with 100mM glucose, was incubated at 37 °C for 1.5 h in 96 well microtiter plate. Non-adherent cells were removed by aspiration with PBS. 200 μL RPMI media with drugs was added to each well and incubated overnight at 37 °C. After incubation, cells were washed with PBS and XTT reagent was added to each well. The plate was incubated in dark for 2 h and colorimetric changes for cell viability were measured at 492 nm using a microtiter plate ELISA reader. The minimum inhibitory concentration for biofilm (MIC_biofilm_) was determined as the concentration that caused a 50% reduction in cell viability.

### Animal studies

The animal experiments were approved by the Institutional Animal Ethics Committee (IAEC) at ICMR-NIRRCH which is recognized by the Committee for the Purpose of Control and Supervision of Experiments on Animals (CPCSEA) under project number 17/21. All the animals recruited for the study were kept at a controlled temperature of 23 ± 1 °C and humidity of 55 ± 5%, with a light cycle of 14 hours on and 10 hours off. The animals had access to water and food ad libitum.

While conducting *in vivo* studies, we carefully considered the ethical guidelines surrounding animal research, specifically the principle of using the minimum number of animals necessary to achieve meaningful results (Festing and Altman, 2002). A positive control, using an antifungal drug with known effects was not included in this study because its efficacy has already been well-documented *in vivo* under similar conditions (Cruz et al. 2018; de Toledo et al. 2020). This study includes both a negative control group and a vehicle control group that allows for rigorous assessment of the drug candidate’s effects while adhering to ethical standards.

### *In vivo* evaluation of besifloxacin for systemic candidiasis

Male C57BL/6 mice (n=17), 8-10 weeks old weighing approximately 25 g were divided into three groups – (i) Control (C), n=5, (ii) Vehicle control (VC), n=6, and (iii) Test group (BS), n=6. An inoculum of 2 × 10^5^ cells of CAL SC5314 was injected intravenously via lateral tail vein in all the groups to initiate systemic infection. Besifloxacin hydrochloride was dissolved in vehicle composed of 20% (v/v) DMSO in normal saline (0.9% NaCl). Drug was administered intravenously 2h post-infection by adjusting the dose to 100 mg/kg/day, which is the NOAEL (No Observed Adverse Effect Level) dose for reproductive toxicity and systemic exposure (Roy et al. 2011). No intervention was performed on the control group post-infection. Filter sterilized vehicle or drug was administered daily for 5 days at the same time. Mice were observed for 5 days for change in body weight and mortality and humanely sacrificed by CO_2_ asphyxiation on day 6^th^. Kidneys were harvested for cfu (colony forming unit) count and histopathology studies.

### Determination of fungal load

#### By agar plate method

For systemic load determination, right kidney of each animal was weighed and macerated in 1 mL sterile PBS. A 1:10 dilution of 100 μL homogenate was plated onto Yeast Extract–Peptone–Dextrose (YPD) agar containing 100 μg/mL chloramphenicol. The plates were incubated at 37 °C for 24-48 hrs. Colonies were counted manually and fungal load was calculated as cfu/organ weight.

#### By histopathology

Dissected kidneys of each animal were fixed in formalin and embedded in paraffin to obtain tissue blocks. Tissue sections of 5 μm were obtained, deparaffinized, and stained using Periodic-Acid Schiff (PAS) and hematoxylin for examination under light microscope. *Candida* load was also measured as percentage of area in the tissue covered by fungal hyphae. Three consecutive readings were obtained from different sites of each tissue section using ImageJ software.

### Statistical analysis

Statistical analysis was performed using GraphPad Prism 8 software. All data were presented as mean ± SD. Non-parametric one-way ANOVA (Kruskal-Wallis test) was used to compare ranks between groups in animal studies. The Synergy Finder plus web tool (Ianevski, Giri, and Aittokallio 2022) was used for statistical interpretation of drug combination assay results.

## RESULTS AND DISCUSSION

Target-specific inhibitors against CAL were shortlisted by identifying evolutionarily conserved proteins in *Candida* that shared homology to drug targets in other pathogens. We initially used 70 target proteins (Chakraborty et al. 2023) corresponding to 35 drugs in Drugbank to identify homologs in *Candida* proteome. Functional domain of DNA topoisomerase IV was found to be conserved among CAL and other bacterial pathogens like *Streptococcus pneumoniae, Haemophilus influenzae* and *Streptococcus pyogenes*.

The fluoroquinolone drugs besifloxacin (BS) and gemifloxacin (GM) target the bacterial DNA topoisomerase IV and DNA gyrase enzymes belonging to the type II topoisomerase family. In bacteria, DNA topoisomerase IV is a heterotetramer of two subunits – subunit A (encoded by *ParC*) and subunit B (encoded by *ParE*), while DNA gyrase consists of two polypeptide subunits, gyrA (homologous to *ParC*) and gyrB (homologous to *ParE*) forming a heterotetramer. In CAL, topoisomerase II is a distant homolog of bacterial topoisomerases IV and DNA gyrase, sharing significant sequence similarity (bit score 493, e-value < 0.0001) (refer to supplementary Table 1). It is a homodimer of a single polypeptide; N- and C-terminal aligns to subunit B (GyrB/ParE) and subunit A (GyrA/ParC) of bacterial topoisomerase IV respectively (Bush Natassja G, Evans-Roberts, and Maxwell 2015) (Figure 1A) Therefore, we postulated that fluoroquinolone drugs may be active against *Candida* by inhibiting topoisomerase II.

**Figure 1.**
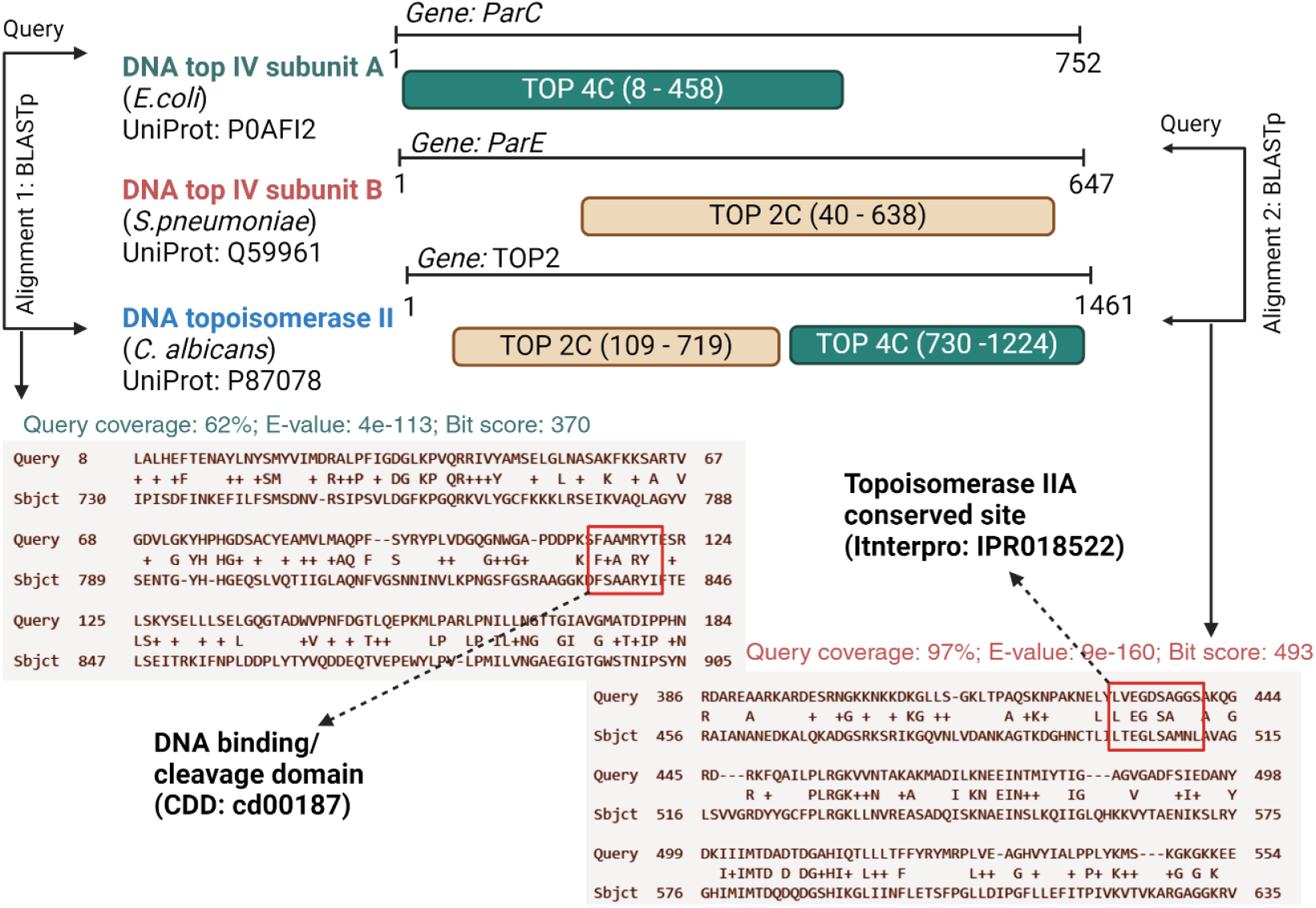
Representative domain architecture and sequence homology between bacterial target of besifloxacin - DNA topoisomerase IV and *Candida* DNA topoisomerase II. All sequences are obtained from UniProt/TrEMBL database. Domain architecture is inferred from InterPro database; alignments are performed using local BLASTp algorithm. Figure created with licensed version of BioRender.

The antifungal potency of BS and GM was determined using *in vitro* broth microdilution assay against *C. albicans*. Inhibition was solely observed at the highest concentration (500 mg/L) of GM, while sustained inhibition was evident across all evaluated concentrations for BS against *C. albicans*. The cell doubling time was calculated from cell concentration data derived from OD600 values as discussed previously (Mukherjee et al. 2021) indicated that BS could impact the growth of CAL comparable to that of fluconazole at higher concentrations ranging from 500 mg/L to 7.8 mg/L. The inhibitory dose (IC_50_) for BS was established at 15.6 mg/L from the above experiment (Figure 2A) and MIC_biofilm_ was found to be 31 mg/L (Figure 2B). GM was not used for further assays due to its poor performance *in vitro*.

**Figure 2.**
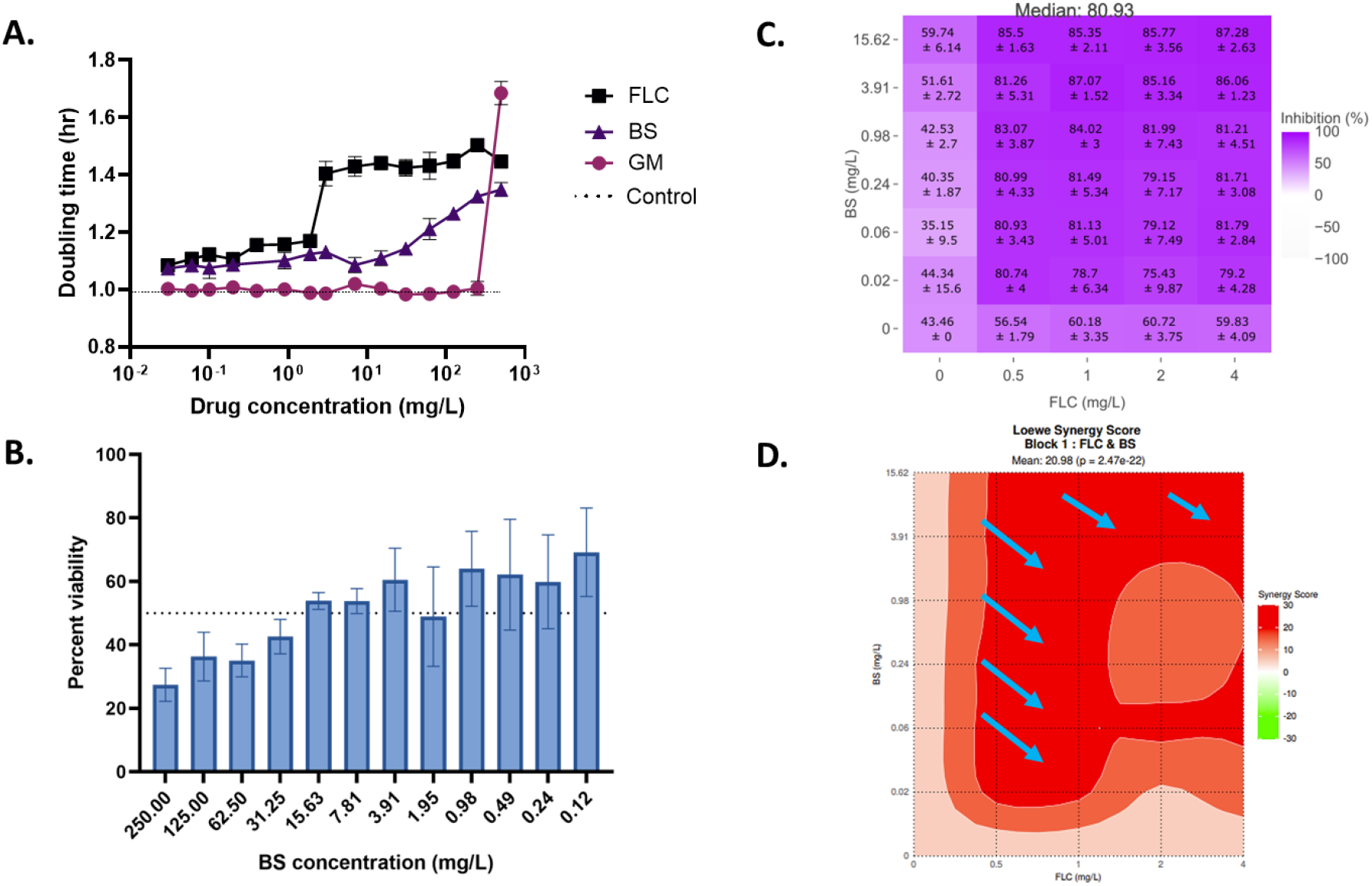
(A) Doubling time graph comparing *C. albicans* growth in terms of normalized doubling time of *C. albicans* calculated using OD_600_ values after 24 h incubation with drugs (individual assay). Doubling time for control is shown as dotted line on y-axis. (B) Anti-biofilm assay of BS. Graph showing % viability of biofilms calculated from OD_490_ values measured using XTT assay. Dotted line marks 50% viability of biofilms. (C) A drug response matrix with mean percentage inhibition for besifloxacin and fluconazole combination assay. (D) A contour plot constructed using Loewe synergy scores derived from the % inhibition of *C. albicans* in the dose response matrix. The areas marked in red and white display regions of synergy and additivity, respectively. Arrows indicate the regions of highest synergy for the combined dose of BS and FLC against *Candida*.

Combination of BS and fluconazole (FLC) reflected a better dose response than the drugs alone. An 80% ± SD inhibition was seen at lowest concentration of BS (0.02 mg/L) and FLC (0.5 mg/L) while 87% ± SD inhibition was seen at the highest concentration (Figure 2C). The Loewe synergy score based on the principle of Loewe Additivity asserts that a drug cannot interact with itself, and the combined effect of two drugs can be determined by their relative potencies (Lederer, Dijkstra, and Heskes 2019). The score is a reflection of synergistic (>10), additive (−10 to 10) and antagonistic (<-10) effects of two or more drug in combinations. Based on this principle, the synergy score (δ = 29.58) for BS and FLC signifies a positive synergistic effect when used in combination suggesting the possibility of clinical utility after further exploration with clinical strains and animal models. We did not observe any areas depicting antagonism (green) in the tested range of the drugs (Figure 2D).

The antifungal effect of BS was further studied in murine model of systemic candidiasis and quantitated in terms of weight change, colony forming unit (cfu) count and histopathology of kidneys. BS has been previously evaluated for systemic and ocular toxicity in rats; no sign of clinical or histopathologic toxicity was observed for doses upto 400 mg/kg/day for 28 days (Roy et al. 2011). Therefore, the NOAEL dose of 100 mg/kg/day was used to evaluate BS against systemic *Candida* infection. A significant reduction in weight was observed for control (C) and vehicle control (VC) groups as compared to drug-treated ones (BS) (Figure 3A). The fungal load in the kidneys of the BS group was significantly lower than that of the control and vehicle control groups (Figure 3B, supplementary figure S1). Histopathological studies showed clearance of fungal hyphae from the kidneys of the treated mice, with the tissue retaining normal histology and a lower number of neutrophils. Fungal hyphae in C and VC groups were accompanied with infiltration of neutrophils indicating infectious inflammation (Figure 3C and 3D). Fungal-infected area in kidney was compared between VC and BS groups using ImageJ software. It was observed that treatment with BS led to an 83% reduction of fungal burden in the treatment group compared to VC (Supplementary table 2, figure S2).

**Figure 3.**
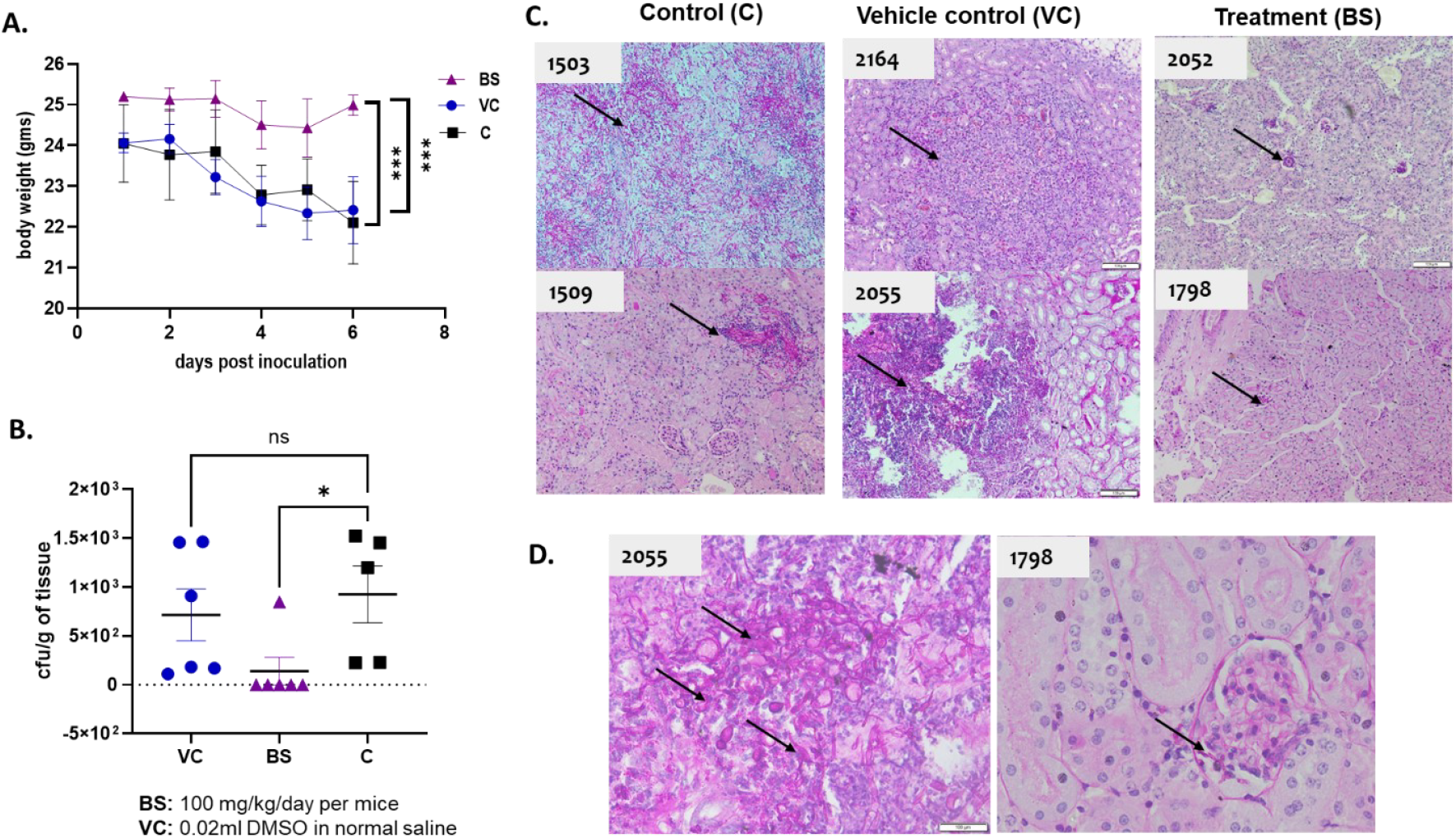
(A) Mean body weight of mice over a period of 5 days post-inoculation. (B) Scatter plot showing viable colony forming units (cfu) per gram of kidney tissue. (C) Histological features of mice kidney stained with PAS showing *C. albicans* infection in C, VC and BS groups at 10X magnification. Arrows indicate fungal hyphae and dark blue dots indicate neutrophil infiltration in tissues. (D) Magnified image (40X) of kidney section from mice in VC and BS groups, respectively. *Candida* yeast and hyphal cells are shown in arrow. Mice identification number is displayed in white box on the upper left corner of images. *** indicates p<0.001, * indicates p<0.01, ns implies not significant.

It has been predicted through *in silico* docking studies that fluoroquinolone antibiotics bind and interact with amino acids ARG841, GLN803, ALA840 within the active site of CAL Topoisomerase II, potentially inhibiting the fungi(Jadhav and Karuppayil 2021).

Furthermore, moxifloxacin and gatifloxacin, which belong to the fluoroquinolone group, have been studied for their *in vitro* antifungal activity against ocular *Candida* species. When used undiluted, these drugs were able to inhibit 95% of fungal growth, demonstrating their antifungal properties (Ozdek et al. 2006). However, the antifungal properties of besifloxacin have not been explored before. The observations from our study confirm the inhibitory activity of the fluoroquinolone, specifically besifloxacin, against *C. albicans*.

### Conclusions

Several efforts are being made globally to identify novel drug combinations that can work synergistically with fluconazole to reduce side effects and overcome resistance (Eldesouky et al. 2021; Li et al. 2019). The observations from our study confirm the inhibitory activity of the fluoroquinolone, specifically besifloxacin, against *C. albicans*. In combination, the MIC reduced to 0.5 mg/L from 2 mg/L (fluconazole alone), showcasing synergistic inhibitory effects on *C. albicans*. Additionally, besifloxacin was observed to prevent the spread of infection in the kidneys of mice. It is important to highlight that, while besifloxacin has FDA approval for human use, utilizing DMSO and saline as a vehicle in this study might not be appropriate for higher primates. Consequently, there is a need to develop a new formulation of this drug specifically for intravenous use. In conclusion, our findings contribute to broadening treatment options for candidiasis, while reducing the risk of antifungal resistance and addressing the challenges of mixed pathogen infections.

## Acknowledgements

The authors are grateful to Dr. Geetanjali Sachdeva, Director, ICMR-NIRRCH for support. This work (RA/1690/06-2024) was supported by was supported by DBT, India [BT/PR40165/BTIS/137/12/2021], SERB, India [CRG/2021/004937] and SRF from ICMR [Myco/Fell/14/2022-ECD-II]. Funding sources were not involved in study design and article preparation.

## Author contributions

SIT, TM and SC participated in the research design; SC conducted *in silico* analysis and experiments; SC, AP and SM conducted the animal studies; SC and AP performed data analysis; SC, AP and SIT wrote the manuscript. All the authors reviewed the manuscript.

