## supplementary Table 1 for "Identification and evaluation of besifloxacin as repurposed antifungal drug in combination with fluconazole against *Candida albicans*"

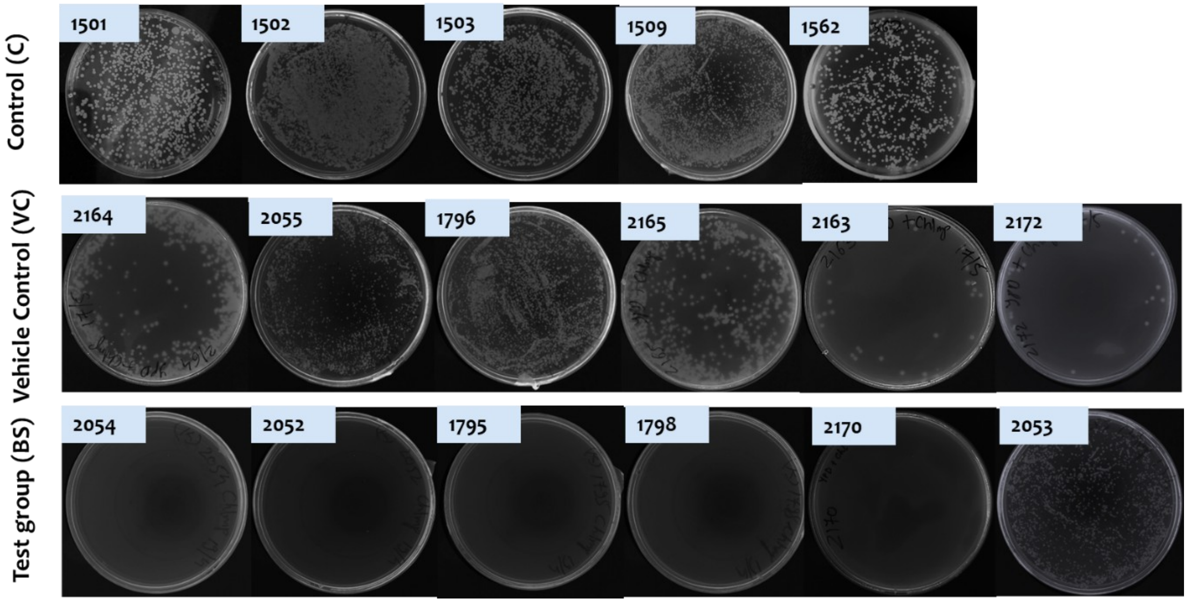


**Supplementary Figure S1.** The kidney fungal burden of *C. albicans* in mice quantified using agar plate assay. Mice identification number is indicated in sky blue boxes on the upper left side of each plate.

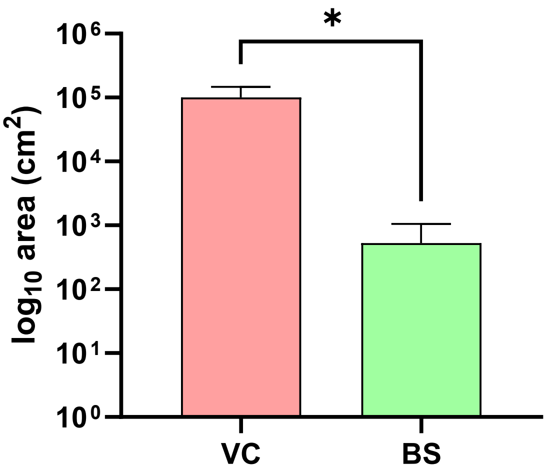


**Supplementary Figure S2.** Fungal infected area between VC and BS group. Infected area is calculated using ImageJ software by demarcating the area filled with fungal hyphae from the non-infected area in photograph of histological sections of kidney taken under the 10X objective. A total of 9 readings from three sections of a single kidney was taken to draw conclusions. Statistical significance ($P < 0.01$) was calculated by unpaired t-test.

| Drug molecule (DrugBank ID) | Target name (UniProt id) | Target homolog in Candida (UniProt id) | Query coverage | E-value | Percent positives | Bit score | Domain description (CDD) | CDD identifier |
|---|---|---|---|---|---|---|---|---|
| Besifloxacin hydrochloride (DB06771), Gemifloxacin (DB01155) | DNA topoisomerase 4 subunit B (Q59961, P20083) | DNA topoisomerase 2 (P87078) | 97% | 9e-160 | 42% | 493 | DNA topoisomerase IV subunit B (Topoisomerase II superfamily) | PRK05559 |
| Besifloxacin hydrochloride (DB06771), Gemifloxacin (DB01155) | DNA gyrase subunit B (P0A4L9, P0AES6) | DNA topoisomerase 2 (P87078) | 97% | 5.71e-132 | 40% | 424 | DNA gyrase subunit B (TopoisomeraseII superfamily) | PRK05644 |
| Besifloxacin hydrochloride (DB06771), Gemifloxacin (DB01155) | DNA topoisomerase 4 subunit A (P0AFI2, P72525, P43702) | DNA topoisomerase 2 (P87078) | 62% | 4e-113 | 40% | 370 | DNA topoisomerase 4 subunit A (Top 4c superfamily) | PRK05561 |
| Besifloxacin hydrochloride (DB06771), Gemifloxacin (DB01155) | DNA gyrase subunit A (P0AES4, P43700, P72524) | DNA topoisomerase 2 (P87078) | 58% | 4.05e-112 | 41.6% | 367 | DNA gyrase subunit A (Top 4c superfamily) | PRK05560 |
| Retapamulin (DB01256) | Large ribosomal subunit protein uL3 (Q9A1X4) | mitochondrial 54S ribosomal protein YmL9 (A0A1D8PRP6) | 98% | 2e-78 | 53% | 238 | 50S ribosomal protein L3 (rpl3p superfamily) | PRK00001 |

**Supplementary Table 1.** List of FDA approved drugs, their targets and corresponding homologous *Candida* proteins identified by delta-BLAST analysis.

| Animal | Total CFUs | Tissue weight in grams | CFUs/gm of tissue | Average CFUs/gm of tissue | Percent Decrease in fungal load |
|---|---|---|---|---|---|
| Vehicle control group (VC) | | | | | 83.23238103% |
| 1796 | 300 | 0.333 | 909.09 | 844.7333333 | |
| 2055 | 300 | 0.338 | 844.73 | | |
| 2163 | 32 | 0.188 | 170.21 | | |
| 2164 | 300 | 0.206 | 1456.31 | | |
| 2165 | 300 | 0.205 | 1463.41 | | |
| 2172 | 28 | 0.154 | 181.81 | | |
| Treatment group (BS) | | | | | |
| 1795 | 0 | 0.215 | 0 | 141.6416667 | |
| 1798 | 0 | 0.212 | 0 | | |
| 2170 | 0 | 0.351 | 0 | | |
| 2052 | 0 | 0.191 | 0 | | |
| 2053 | 300 | 0.353 | 849.85 | | |
| 2054 | 0 | 0.182 | 0 | | |

**Supplementary Table 2.** Percent decrease in fungal load of Besifloxacin treated mice compared to the vehicle control group. (CFU: Colony Forming Unit)
Percent Decrease in fungal load = {[Average CFUs/gm of tissue (VC) - Average CFUs/gm of tissue (BS)]/Average CFUs/gm of tissue(VC)} * 100

(Ref: Lynn Miesel, Kun-Yuan Lin, Voon Ong “Rezafungin treatment in mouse models of invasive candidiasis and aspergillosis: Insights on the PK/PD pharmacometrics of Rezafungin efficacy”)
